# Visualization of the microstructure and distribution of follicles in deep human ovaries using speckle-modulated optical coherence microscopy

**DOI:** 10.1101/2025.08.20.671419

**Authors:** Koichiro Ito, Momoko Kanaya, Seido Takae, Nao Suzuki, Kosuke Tsukada

## Abstract

Ovarian tissue cryopreservation (OTC) and transplantation (OTT) are the only fertility preservation (FP) options for children, adolescents, and young adults (CAYA) who don’t enough time until treatment of primary disease. OTC is still unestablished FP options, but it is expected that selective preservation of ovarian tissue with high follicle density will enhance pregnancy efficiency. Despite its value as a noninvasive follicle visualization method, optical coherence microscopy (OCM) lacks sufficient spatial resolution, imaging depth, and noise suppression capabilities to resolve fine follicular structures in deep ovarian tissue. In this study, we applied speckle modulation to a wavelength-swept OCM optical system in the near-infrared region to visualize the fine structure of follicles distributed in deep tissues. OCM images of 4-day-old mice revealed dense superficial-layer distributions of primordial follicles. These follicles were distinguishable from primary follicles, which were surrounded by a single layer of granulosa cells. In 14-day-old ovarian tissue, secondary follicles exhibited spatial dominance, and the fine structures constituting the follicles were successfully visualized. In ovarian tissues extracted from a patient diagnosed with acute myeloid leukemia (AML), the fine structures of primordial follicles were visualized to a depth of 300 μm, and a comparison with HE staining revealed that follicle density varied significantly depending on the ovarian regions. Speckle modulation-assisted OCM visualized the microstructure of follicles with varying degrees of maturation in human ovaries at the depth necessary for clinical application. Overall, this technology enables quantitative ovarian reserve assessment and selective cryopreservation of ovarian tissue rich in immature follicles, supporting FP in CAYA diagnosed with cancer.

## 1. Introduction

Radiation and chemotherapy for cancer treatment in women of the children, adolescents, and young adults (CAYA) generation often result in ovarian dysfunction or loss [1,2]. The decline in childhood cancer mortality in recent years has increased interest in post-remission quality of life [3]. In this context, ovarian tissue cryopreservation (OTC) technology is emerging as a potential fertility preservation (FP) therapy for the next generation.

In OTC, clinicians extract ovarian tissue from individuals diagnosed with cancer before chemotherapy or radiation and preserve it in liquid nitrogen for long-term storage. The tissues are then autotransplanted following the patient’s desire for pregnancy, thereby preserving fertility [4–7]. Since the initial report in 2004 [8], 130 cases were reported in 2017 [9], and approximately 200 cases were estimated to have been reported by 2020 [10]. Most OTTs have resulted in adequate restoration of ovarian function [11,12]. The functional lifespan of transplanted ovarian tissue depends on its primordial follicle count [13]. Furthermore, follicles within ovarian tissue exhibit heterogeneous distribution, with primordial follicle density varying by two or more orders of magnitude within a single cortical section [14]. Consequently, the primordial follicle density of transplanted ovarian tissue governs transplantation efficacy. Establishing a noninvasive method to visualize and quantify follicle counts holds clinical significance.

Follicles consist of oocytes surrounded by granulosa and theca cell layers. As these follicles mature, their microstructure undergoes characteristic changes. For instance, the granulosa cells surrounding the oocyte exhibit a flattened morphology in primordial follicles, transition to a cuboidal configuration in primary follicles, and subsequently undergo a multilayering process to mature into secondary follicles. Primordial follicles—with an approximate diameter of 20–30 μm—are relatively abundant in the ovarian cortex at a depth of 100–1,000 µm [15]. Therefore, visualizing follicular microstructure, identifying maturation stages, and quantifying density require an imaging depth of at least 100 µm and spatial resolution of approximately 10 µm.

Optical coherence tomography (OCT) is a noninvasive imaging technique that visualizes the internal structure of tissues to a depth of several millimeters [16] and is applied in ophthalmology [17,18], dentistry [19,20], gastrointestinal tract [21,22], coronary vessels [23,24], colon [25], and breast [26]. Recent studies have reported OCT utilization for imaging ovarian tissue [27–29]. Primordial follicles in mouse, bovine, and human ovaries can now be visualized using full-field OCT (FF-OCT) with a spatial resolution of approximately 1 µm [30–32]. We have previously demonstrated the feasibility of evaluating the localization of primordial follicles and ovarian reserve in human ovaries [31]. However, the limited imaging depth (∼100 µm from the tissue surface) and speckle noise, which obscures follicular microstructure, hinder clinical application.

Researchers have proposed speckle noise reduction methodologies from optical and image processing perspectives to recover fine structural information. Those for mitigating optical speckle noise include angular compounding [33], averaging algorithm [34], frequency compounding [35], and polarization diversity [36]. However, these methods compromise resolution and are incapable of eradicating speckle noise. Conversely, adaptive filters, wavelet analysis [37–41], and machine learning techniques can be utilized to eliminate noise from OCT images [42–46]. Although several methods exist for noise reduction in OCT images of the retina [34,47,48], methods suitable for the ovary, which has more structural dispersion than the retina, remain unestablished. Gaussian filters proved effective for noise reduction in segmented OCT images of mouse ovaries [49,50]. Image processing methods such as filtering reduce speckle noise, but accurately restoring noise-degraded structural information without compromising resolution remains fundamentally challenging.

Recently, Liba et al. proposed speckle modulation—which enables the creation of an unlimited number of uncorrelated speckle patterns by moving a diffuser plate within the optical path—and demonstrated that speckle noise can be effectively reduced without degrading resolution [50]. Beyond preserving resolution and reducing noise, the implementation simplicity of speckle modulation in OCT systems supports its suitability for visualizing follicular microstructure.

This study introduces an optical coherence microscopy (OCM) system capable of visualizing follicular microstructure across maturation stages at depths with high follicle density. Specifically, applying phase shift–based speckle noise reduction processing to an optical system based on a 1,300-nm wavelength-swept OCM enabled visualization of follicular microstructures at clinically relevant depths. Using OCM images, we demonstrated, for the first time, the spatial heterogeneity of human primordial follicle density, highlighting the potential for clinical application in FP for CAYA diagnosed with cancer.

## 2. Materials and Methods

### 2.1 Setting up the OCM system

Figure 1 shows an OCM system for noninvasive visualization of follicular microstructure. A swept-source laser (HSL2100, Santec) with a 1,310 nm center wavelength, 171 nm bandwidth, and 20 kHz frequency served as the light source. The laser light passing through the circulator (CIR1310-APC, Thorlabs) was split into reference and sample arms in a 50:50 ratio using a fiber coupler (TW1300R5A2, Thorlabs). The reference light was reflected by a mirror via a polarization controller. The sample light was irradiated onto the sample via a galvanometer scanner, a diffusion plate (DG10-600, Thorlabs) mounted on a linear motion stage, and a 20× water-immersion objective lens (XLUMPLFLN20XW, Olympus) with a 1.0 aperture number. A differential amplifier (PDB570C, Thorlabs) detected the interference signal between the sample backscatter and reference mirror reflection. Data acquisition and processing were performed using a DAQ board (HAD-5200B-S, Santec) and original software.

**Figure 1.**
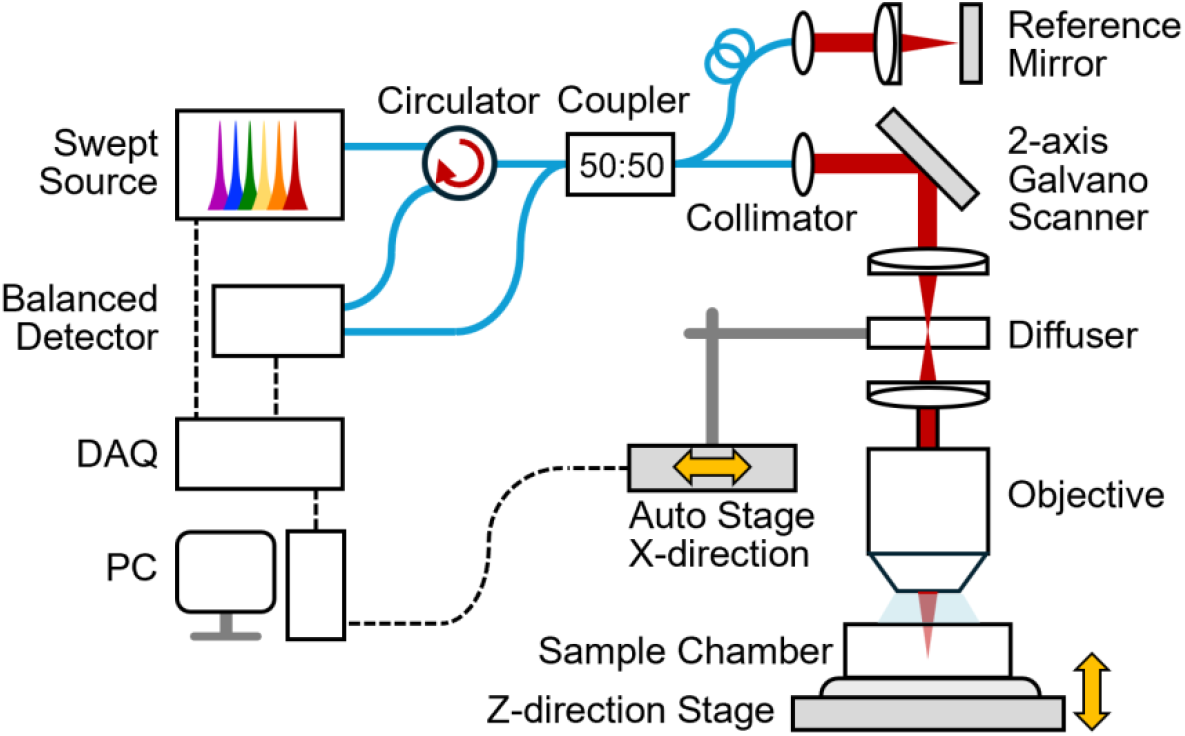
Schematic of the OCM optical system. The laser light from the sample arm is reflected by a galvo scanner and irradiated onto the ovary via a diffuser used for speckle modulation and an objective lens.

A USAF 1951 contrast resolution target was placed on the objective lens’ focal plane, and the lateral spatial resolution was evaluated from the OCM images obtained. Depth resolution was evaluated as the full width at half maximum (FWHM) of the point spread function (PSF), averaged over 20 A-scans acquired with a mirror positioned at the focal plane.

### 2.2 Preparation of the ovarian phantom

An ovarian phantom containing glass beads embedded in polyacrylamide gel was prepared. Specifically, 4.9% polyacrylamide gel was mixed with Intralipid at a concentration of 5% as a scattering agent [51,52], and glass beads with a diameter of 30 ± 2 µm (SPL-30, UNITIKA), mimicking ovarian follicles, were suspended and gelified. The phantom was then placed in a custom-designed sample holder filled with double-distilled water.

### 2.3 Mouse and human ovarian tissue samples

Animal experiments were performed according to protocols approved by the Animal Experiment Committee of the Graduate School of Medicine, St. Marianna University (Protocol Nos. #2304008, #2304009). Ovarian tissues were collected from female mice at 4 days, 14 days, and 20–24 weeks of age after sacrifice through cervical dislocation and were placed on ice. The average body weights of the mice were 3.4 g (day 4, n = 3), 7.9 g (day 14, n = 3), and 39.7 g (week 20–24, n = 3), respectively. Within 12 h of collection, the tissue was placed in a sample chamber filled with phosphate buffer saline, and OCM images were acquired.

Human tissue experiments were approved by the Clinical Trials Subcommittee of the Institutional Review Board for Biomedical Research at St. Marianna University School of Medicine (Approval No. 6272, IN000023141). The details of this clinical study were explained to all patients or their families by their study investigators, and written informed consent was obtained. This study was conducted in compliance with the Declaration of Helsinki (revised in 2024), “Ethical Principles for Medical Research Involving Human Participants, “Ethical Guidelines for Epidemiological Research”, “Nuremberg Code,” and “Act on the Protection of Personal Information”. Ovarian tissue was obtained from a female patient with acute myeloid leukemia (AML) who developed secondary to familial thrombocytopenia at the age of 15 years. The cryopreserved ovarian tissue, approximately 10 mm square and 1 mm thick, was thawed and cut into four equal pieces, each approximately 5 mm square. The tissue was placed in a sample chamber filled with culture medium (G-MOPS™ Vitrolife, Gothenburg), and OCM imaging was performed.

### 2.4 HE staining

The ovarian tissues used for OCM imaging were fixed in 4% paraformaldehyde. They were embedded in paraffin wax, sectioned (5 μm thick), and stained with hematoxylin and eosin following standard protocols.

### 2.5 Process for acquiring OCM images

The laser was irradiated vertically onto the specimen through an objective lens to obtain OCM signals in the depth direction (A-scan). By repeatedly scanning the laser horizontally with a galvo mirror (B-scan), volume data with an area of 666 × 666 µm^2^ and a depth of 75 µm was acquired. The operation of moving the diffusion plate in the B-scan direction at a speed of 2.4 µm/s for 50 µm was repeated 20 times, and volume data was acquired each time and then averaged. En-face images were generated from these data and smoothed with a Gaussian filter (standard deviation 0.8). To compensate for the shallow depth of field of the high-numerical-aperture-objective lens, the lens was axially shifted in 30 µm increments to repeatedly acquire volumetric data. All data were combined into a single 3D image after removing overlapping parts. Three-dimensional images of phantoms and mouse ovarian tissues were generated using the open-source software 3D Slicer, with glass beads and oocytes marked as spheres.

## 3 Results

### 3.1 Evaluation of lateral and depth resolution

The OCM image of the 1951 USAF test target shown in Figure 2(a) was spatially resolved down to the smallest element in the horizontal and vertical directions, indicating that the horizontal spatial resolution was < 4.38 µm. The theoretical value calculated from the FWHM of the wavelength spectrum of the light source was 5.78 µm in air. The discrepancy between the measured and theoretical values resulted from deviations of the wavelength spectrum from an ideal Gaussian profile. Figure 2(b) shows the PSF in the depth direction obtained by placing a mirror on the objective lens’ focal plane, and the depth resolution in air was read as 8.83 µm. The *in vivo* spatial resolution, calculated using water’s refractive index (n = 1.33), was 3.29 µm laterally and 6.64 µm axially, enabling 3D visualization of 30-µm diameter primary follicles and fine structures of secondary follicles.

**Figure 2.**
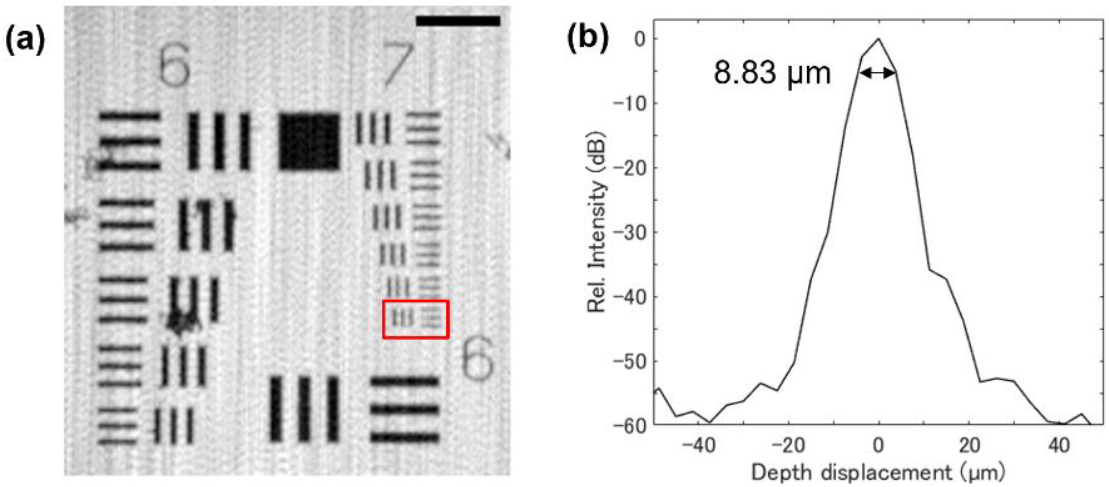
Evaluation of spatial resolution in the horizontal and axial directions. (a) OCM image of the 1951 USAF resolution test chart. The resolution indicated by the red frame is less than 4.38 µm. (b) Point spread function obtained in the depth direction by the A-scan. The depth resolution in air is 8.83 µm.

### 3.2 Ovarian phantom imaging test

Glass beads (diameter: 30 μm) were visualized at a depth of at least 700 μm from the ovarian phantom surface (Figure 3(a)). The clear shadows of the black circles (white arrows) indicate the actual beads, while the blurred shadows (red arrows) indicate the signal attenuation caused by the beads above. The white spots in the center of the shadows are light reflections caused by significant refractive index changes at the top and bottom of the beads (yellow arrows). Figure 3(b) and S1 Video show the cross-sectional images in the B-scan direction. In the deeper regions, the signal contrast with the surrounding area decreased compared to the surface region due to light attenuation caused by scattering and absorption. Figure 3(c) and S2 Video 2 show the distribution of glass beads detected in a volume of 666 × 666 × 810 µm^3^. These results suggest the feasibility of visualizing 30 µm-diameter primordial follicle distributions in deeper ovarian regions.

**Figure 3.**
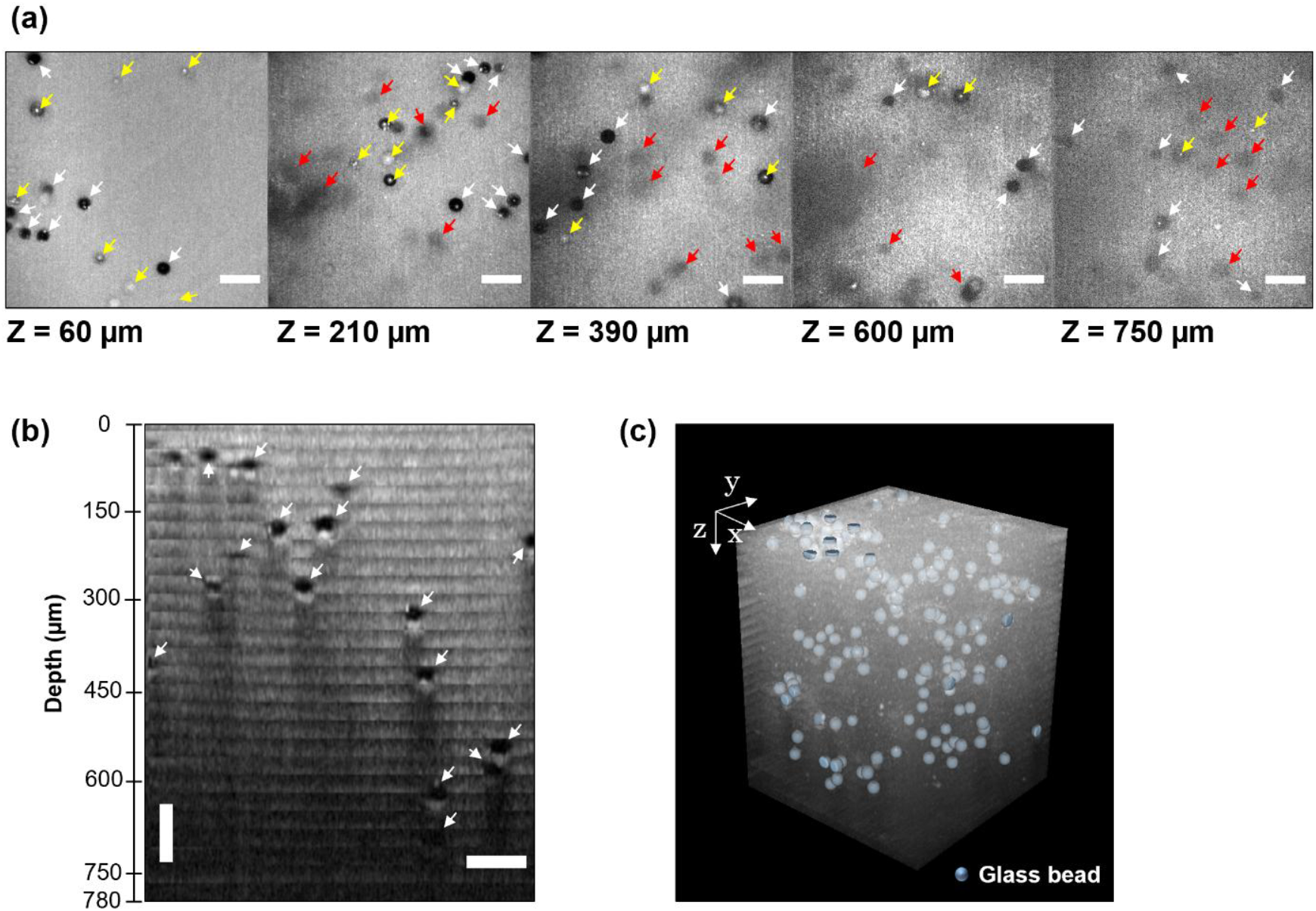
OCM images of an ovarian phantom. (a) En-face images at different depths (Z = 60–750 µm) from the surface. White arrows indicate the actual beads present at the focal plane, while red arrows indicate shadows caused by signal attenuation due to beads in the upper layers. The white dots shown by the yellow arrows represent strong light reflection caused by refractive index changes at the upper and lower surfaces of the beads. (b) Typical XZ cross-sectional (b-scan) image. Refer to S1 Video. (c) 3D diagram of the glass bead distribution. Refer to S2 Video. The scale bar indicates 100 µm.

### 3.3 Visualization of follicles in mouse ovaries

Figure 4 shows OCM images of mouse ovaries at 4 days (a), 14 days (b), and 20–24 weeks (g, h) of age, reaching a 300-µm depth. In 4-day-old mouse ovaries, primordial follicles with diameters of approximately 20–40 µm appeared as dark circular shadows and were localized relatively close to the tissue surface. With increasing depth, primary follicles were also occasionally observed (Figure 4(a)-1, -2, S3 Video). For example, Figure 4(c), an enlarged view of the yellow dashed box in Figure 4(a)-3, shows primary follicles surrounded by a single granulosa cell layer (yellow arrows). In the HE-stained image of the same ovarian tissue shown in Figure 4(d), primary follicles covered by a single granulosa cell layer (yellow arrows) were also observed, suggesting that the fine structures visualized by OCM were appropriate. The density and localization of primordial and primary follicles varied among individuals even at the same depth.

**Figure 4.**
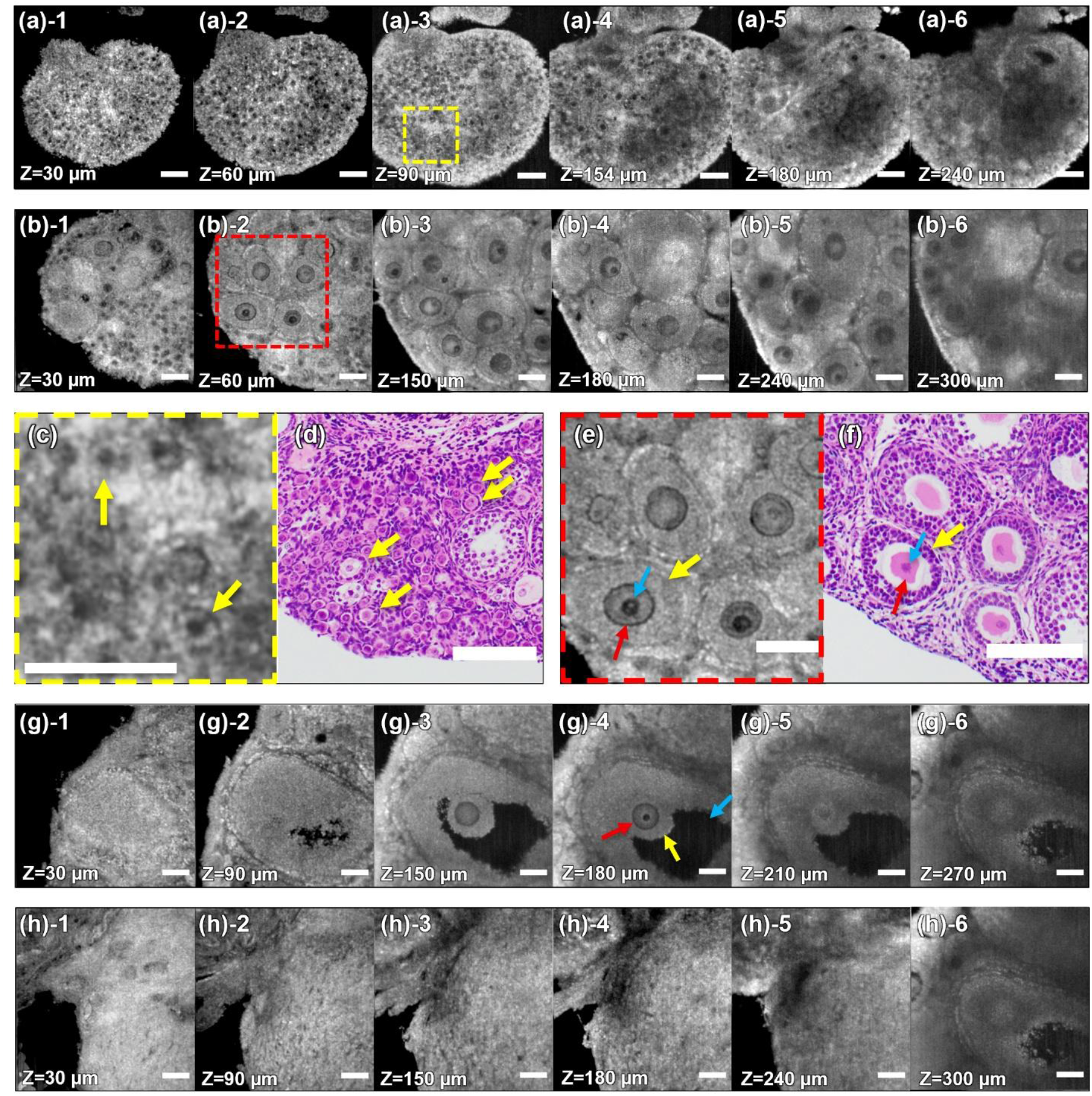
OCM images of mouse ovaries. (a) En-face images at depth Z from the tissue surface of 4-day-old (a) and 14-day-old (b) mice. Refer to S3 and S4 Videos, respectively. (c) and (e) are enlarged views of the dotted square areas in (a)-3 and (b)-2, respectively. (d) and (f) are HE-stained tissue sections of the same tissue used for (c) and (e). (g) and (h) are en-face images at depth Z from the tissue surface of mouse ovaries at 20–24 weeks of age. Refer to S5 Video. All scale bars are 100 µm.

In 14-day-old mouse ovaries, primordial and primary follicle counts decreased compared to those at day 4 and, as shown in Figures 4(b)-1 and -2, were localized within ∼100 µm of the tissue surface. Beyond 100 µm depth, secondary follicles and cyst-like follicles (100–300 µm in diameter) dominated the tissue, with larger follicles concentrated in central regions (Figure 4(b)-3 to -6, S4 Video). In Figure 4(e), an enlarged view of the red-framed area in Figure 4(b)-2, the oocytes (red arrows) and their nuclei (blue arrows) of the secondary follicles—and the thickened granulosa cells (yellow arrows)—were clearly visualized, corresponding to the HE image of the same tissue shown in Figure 4(f).

In the ovaries of 20- to 24-week-old mice shown in Figure 4(g), follicles ≥ 600 µm in diameter were observed. In addition to secondary follicle structures, the oocyte (red arrow), cumulus cells (yellow arrow), and follicular cavity (blue arrow) were clearly visualized ((g)-4, S5 Video). In addition, areas with almost no follicles were identified, as shown in Figure 4(h).

Figure 5 shows the 3D distribution of follicles based on the OCM images shown in Figures 4(a) and 4(b). The slice interval of the OCM images was 3.75 µm, and follicles overlapping multiple slices were counted as one. Primordial/primary follicles and secondary follicles were continuously observed in approximately 5–13 and 14–50 layers, respectively. In 4-day-old mouse ovaries, primordial/primary follicles are predominantly located in the peripheral regions of the ovary, while secondary follicles are distributed in the central region, as shown in the 3D images (Figure 5(a), S6 Video). Similarly, in 14-day-old mouse ovaries, primordial and primary follicle density was lower than in 4-day-old ovaries, with follicles scattered throughout the peripheral regions (Figure 5(b), S7 Video).

**Figure 5.**
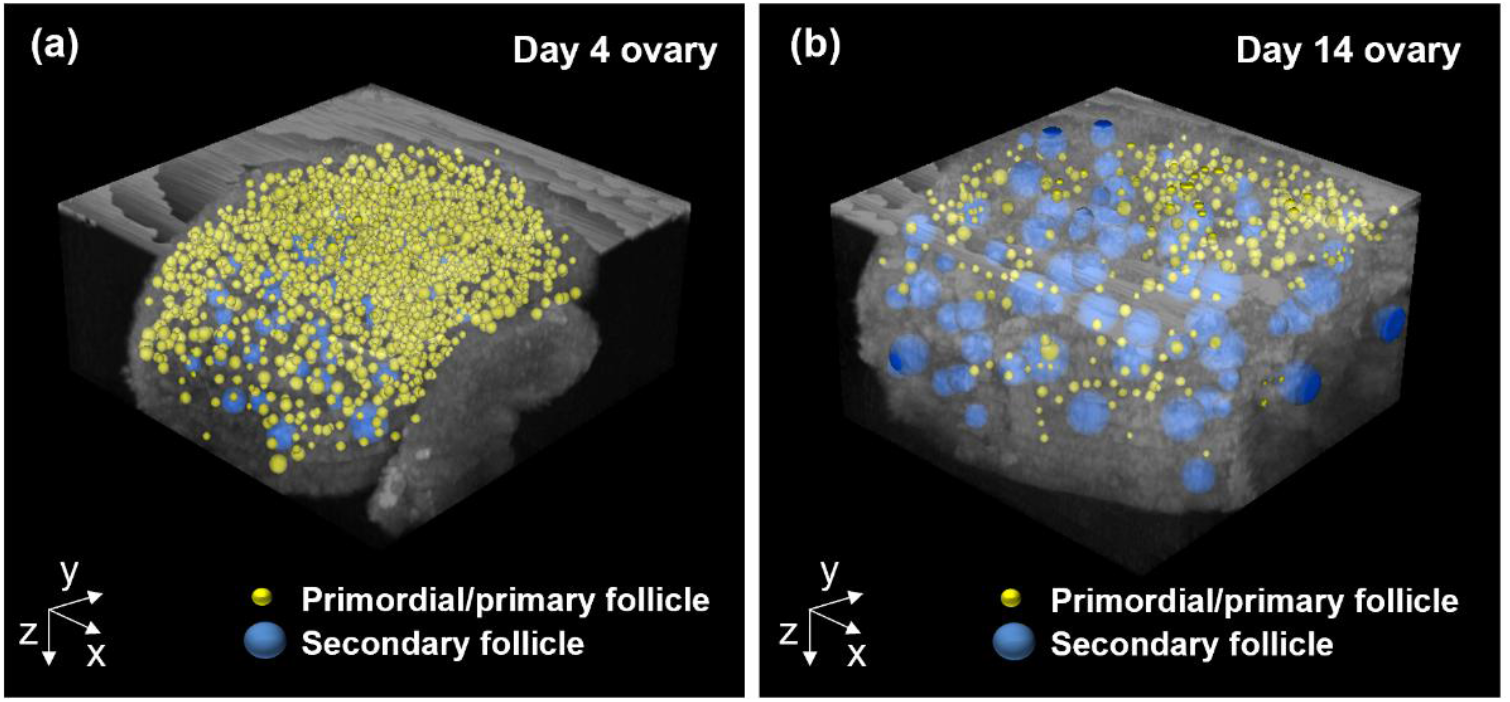
(a) Three-dimensional distribution of follicles in mouse ovaries at 4 days of age and (b) at 14 days of age. Yellow and blue spheres indicate primordial/primary follicles and secondary follicles, respectively. Refer to S 6 and S7 Vidoes.

### 3.4 Localization imaging of human follicles

Figure 6(a) shows an en-face OCM image of ovarian tissue from a 15-year-old AML patient at a depth of 120 μm from the surface. A group of primordial follicles was observed in the parenchymal tissue (upper left half) and distributed in the spindle-shaped interstitial tissue (lower right half). In Figure 6(b)—an enlarged view of the yellow dashed box in Figure 6(a)—the primordial follicles in the OCM images were clearly visualized, including the oocytes (yellow arrows) and their nuclei (red arrows). Their structure was consistent with the HE-stained images of the same ovarian tissue shown in Figure 6(c). The theca cell layer surrounding the primordial and primary follicles was not discernible in HE-stained images. In the OCM images, mouse primordial follicles appeared as dark shadows, while human primordial follicles resembled structures observed in HE-stained sections. The mouse ovaries were imaged under sustained physiological conditions postovariectomy, whereas the human ovaries were observed postcryopreservation. The optical properties of the follicles, specifically differences in tissue or cell water content, refractive index, and scattering coefficient, may have caused the differences in OCM images.

**Figure 6.**
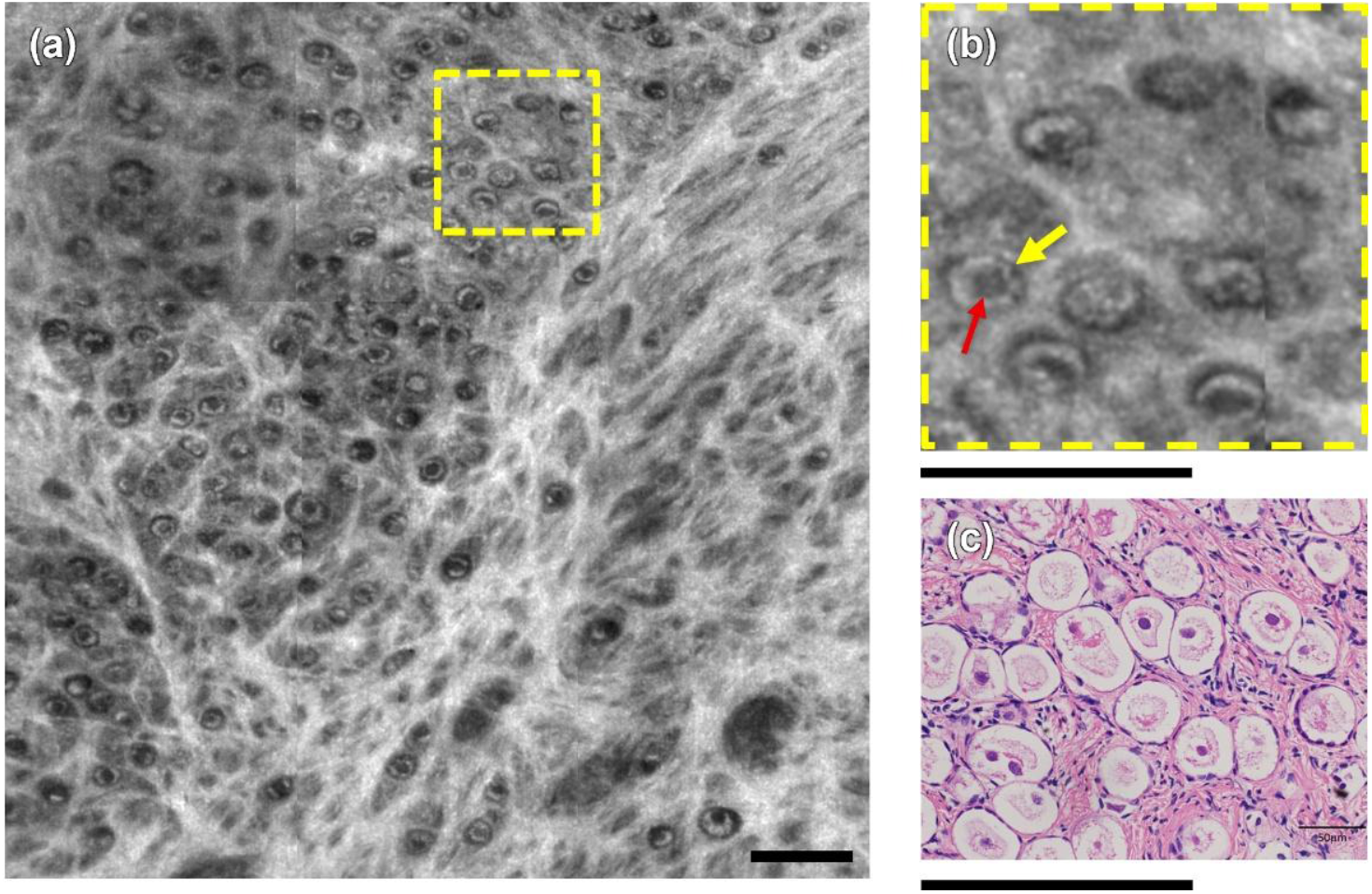
(a) Frontal OCT image at a depth of 120 µm from the surface of the ovary of a 15-year-old female patient with acute myeloid leukemia. A cluster of primordial/primary follicles is observed in the cortex on the upper left side, and primordial follicles are also distributed in the ovarian stroma rich in fibroblast-like spindle cells on the lower right side. (b) Magnified view of the yellow dotted rectangle in (a), with yellow and red arrows indicating oocytes and their nuclei, respectively. No capsule formation is recognized around the primordial/primary follicles. (c) HE-stained sections of the same tissue. The circular structures that clearly show cellular and nuclear-like structures are considered to be primordial/primary follicles. The scale bar corresponds to 200 µm.

Figure 7 shows OCM images from the surface to a depth of 300 μm at different regions of interest (ROI (a) to (c)) on human ovarian tissue fragments (S8 Video). Follicle density and diameter varied with ROI and imaging depth. For example, in ROI (a), follicles were distributed from the surface to a depth of 180 μm, whereas in ROI (b), they were concentrated in the region from 120 to 180 μm in depth. In contrast, in ROI (c), follicles were widely distributed from 120 to 300 μm in depth. Regional differences in follicle density are also evident in the HE-stained cross-sectional images shown in Figure 7(d), which clearly reveal areas of high follicle density alongside regions nearly devoid of follicles. This suggests the importance of selective cryopreservation of ovarian tissue with a higher follicle density.

**Figure 7.**
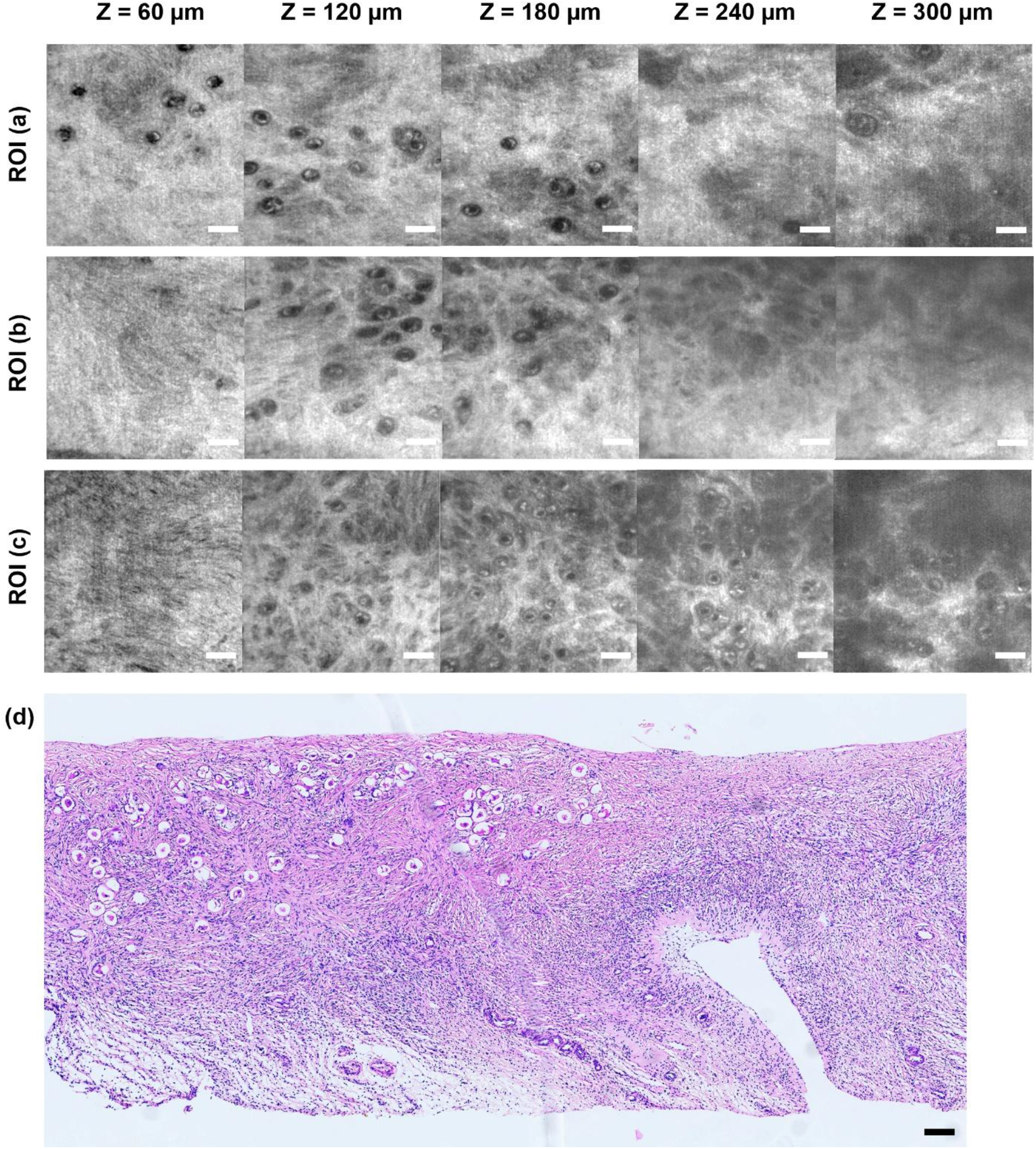
In the same ovary as in Figure 6, an uneven distribution of primordial follicles was observed in the regions of interest (a) to (c) and at depths of 60–300 µm. (d) A low-magnification HE-stained image of the same ovarian tissue similarly showed a region-dependent distribution of follicles. The scale bar corresponds to 100 µm. Refer to S8 Video.

## 4. Discussion

### 4.1 Visualization of deep follicular microstructure

The newly developed near-infrared OCM achieved two key advances: visualization of primordial follicular microstructure at depths up to 300 µm in mouse and human ovarian tissue, and imaging of follicle density distribution in human ovaries using a clinically adaptable light irradiation method.

Quantifying follicle localization and density by the maturation stage is essential for evaluating ovarian reserve; thus, visualizing deep follicular microstructure marks a significant advance toward clinical application. Previous OCT-based follicle imaging was limited by insufficient spatial resolution and noise removal, making it difficult to distinguish between primordial and primary follicles. As shown in Figure 4(c) and S3 Video, we visualized a single layer of granulosa cells in the primary follicle. However, our setup was limited to a depth of 300 μm, whereas primordial/primary follicles exist at a depth of approximately 1 mm from the ovarian surface. Therefore, further expansion of the imaging depth is desirable to quantify the total number of follicles in the tissue. Lasers with a wavelength band of 1,700 nm are suitable for imaging deeper tissue due to their high light penetration [53–55]. Although the extension of the wavelength band reduces spatial resolution, the combination with speckle modulation may allow deep imaging while maintaining the required spatial resolution.

We noninvasively visualized the heterogeneous distribution of human primordial follicles. As shown in Figure 6, follicle density was nonuniform, with primordial follicles present in the parenchymal and interstitial tissue. As shown in Figure 7, we demonstrated that follicle density varies with both lateral position relative to the tissue surface and imaging depth. These results align with HE-stained images from the same patient tissue and support previous findings that follicle distribution is nonuniform, with density varying by more than two orders of magnitude between sections of the same ovary [14]. Selective cryopreservation of tissues with high follicular density may directly improve treatment outcomes. Conversely, freezing ovarian tissue that contains almost no follicles poses risks to patients in terms of cost and loss of fertility.

### 4.2 Accelerating OCM imaging for clinical use

In general, culturing oocytes within the ovary and maintaining their viability is challenging and requires careful attention to temperature, gas partial pressures (5–6% O_2_ and CO_2_ in vivo), and shielding (oocytes are light sensitive). Therefore, maintaining follicular viability while rapidly quantifying density and localization in human ovarian tissue requires faster OCM image acquisition to minimize examination time in clinical settings. Acquiring a volumetric dataset with a 666 µm lateral width and 75 µm depth required approximately 3 min. Additionally, utilizing high-aperture objective lenses necessitated repeated imaging at multiple focal depths, further increasing the acquisition time.

Speckle modulation was key to visualizing the follicular microstructure. As shown in Figure 8, the fine structures of mouse primordial follicles (a, b) and secondary follicles (d, e)—and human primordial follicles (g, h)—were not visible in the original or Gaussian-filtered images. However, they were clearly visualized only when speckle modulation and Gaussian filtering were combined (c, f, i). As described in the method, applying speckle modulation requires acquiring 20 volumetric datasets via laser irradiation through a scattering plate—a bottleneck that limits acquisition speed. Implementing machine learning algorithms offers a promising approach to reducing the number of required data acquisitions [56]. The precision and accuracy of object detection are contingent on learning using teacher images with clearly identified correct answers. Consequently, machine learning algorithms trained on follicle images accurately identified via speckle modulation are expected to enable high-precision follicle detection, even from low-quality images with limited averaging.

**Figure 8.**
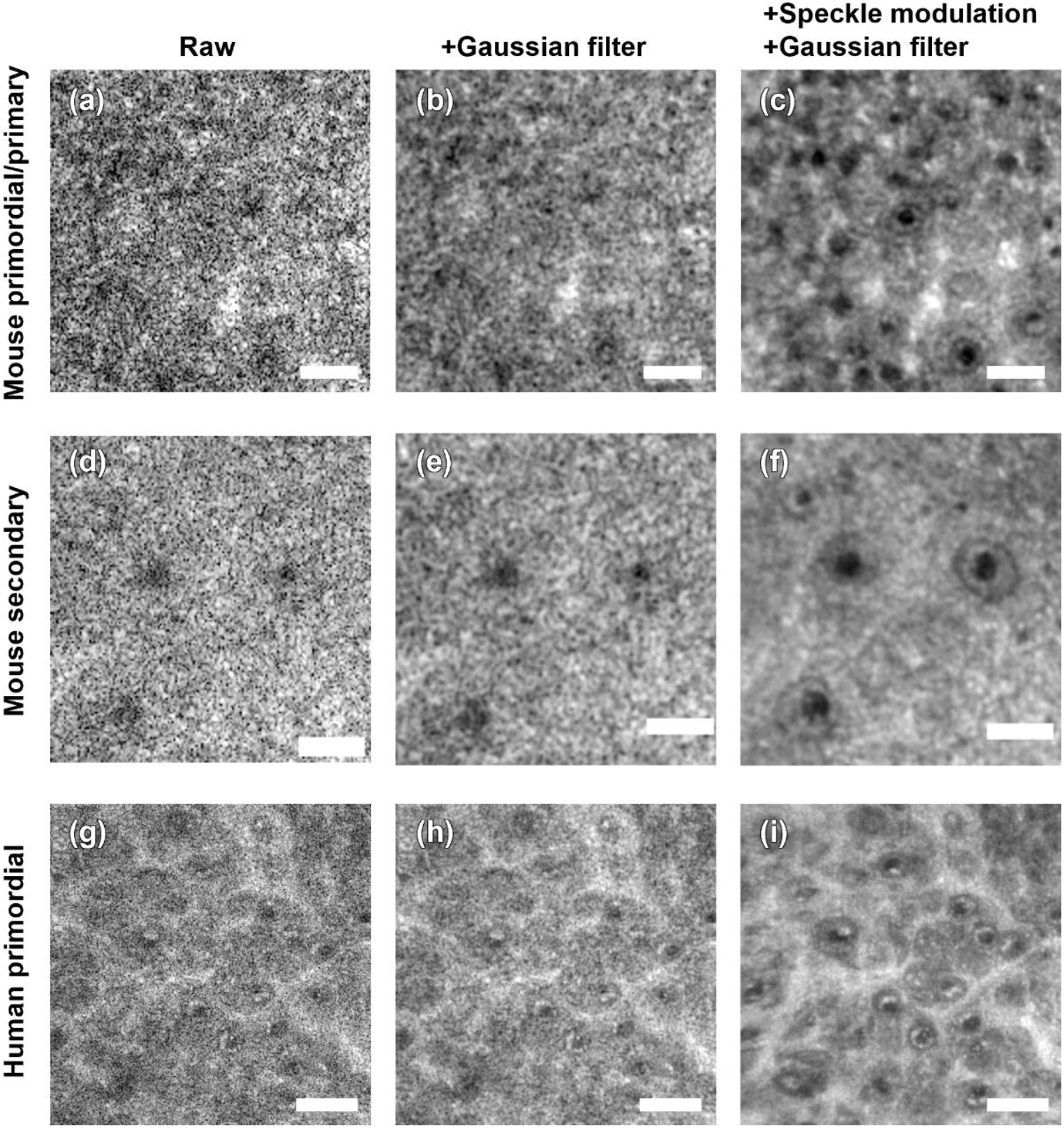
Typical examples of noise reduction using speckle modulation to visualize the fine structure of follicles. OCM raw images containing speckles of mouse ovaries of (a) mouse primary/primary follicles, (d) secondary follicles, and (g) human ovarian tissue. (b, e, h) Images processed with a Gaussian filter applied to the original images, and (a, d, g) effects of speckle reduction processing using a diffusion plate. Scale bars: 50 µm for mouse ovary images (a–f) and 100 µm for human ovary images (g–i).

Using a high-speed light source also effectively reduces acquisition time. This study employed a 20 kHz source; however, recent reports have demonstrated light sources operating at 325 kHz [57], which is expected to reduce image acquisition time.

### 4.3 Assessment of possible risks and safety

Presently, OTC is considered a promising FP strategy for children and adolescents diagnosed with cancer who face chemotherapy or radiation therapy that may impair ovarian reserve [58]. OTC has been clinically applied in Europe and the United States since the late 1990s. With increasing reports of live births in recent years, it is no longer considered an experimental technology in these regions [59,60]. For CAYA cancer patients who cannot delay treatment, OTC remains the only FP option. Consequently, clinical research on OTC safety and efficacy should proceed in parallel with the development of follicle visualization technology. A previous study demonstrated that in vitro fertilization using OCT-imaged mouse ovaries did not reduce oocyte yield, fertilization rate, or blastocyst formation compared to controls, and no increase in congenital anomalies was observed in offspring [32]. The wavelength, irradiance, irradiation time, energy, and heat generated required for follicle visualization depend on the OCT setup, so the noninvasiveness of the method for germ cells must be re-evaluated. Increasing irradiation energy or scan count improves image quality and raises the risk of tissue damage. Defining the damage threshold from laser exposure to oocytes using primate and human ovarian tissue is essential to ensure safe transplantation.

Furthermore, in patients diagnosed with leukemia (the most common childhood cancer), neuroblastoma, Burkitt lymphoma, ovarian cancer, and related conditions, frozen-preserved ovarian tissue may carry a risk of minimal residual disease (MRD), potentially precluding transplantation. OCT proves effective for tumor detection [61–66]. A future OCT-based technique capable of simultaneously visualizing follicular density and detecting MRD lesions may benefit patients for whom ovarian tissue transplantation is not feasible.

## 5. Conclusion

This study presents the first noninvasive visualization of follicular microstructure by combining speckle noise reduction via a diffusion plate with 1,300-nm wavelength OCM. Differentiating primordial, primary, and secondary follicles at a 300 µm depth in human ovarian tissue enables high-accuracy quantitative assessment of ovarian reserve. By reducing imaging time and verifying safety tests for laser irradiation, this technology is expected to contribute to accelerating OTC and transplantation treatment for patients with CAYA-generation cancer.

## Supporting information

S1 Video: Longitudinal section (B-scan) imaging of an ovarian tissue phantom

S2 Video: Three-dimensional distribution of glass beads in ovarian tissue phantom

S3 Video: Imaging of primordial follicles in the ovaries of 4-day-old mice

S4 Video: Secondary follicle imaging of 14-day-old mouse ovaries

S5 Video: Visualization of mature secondary follicles with follicular cavities

S6 Video: 3D distribution of follicles in the ovaries of 4-day-old mice

S7 Video: 3D distribution of follicles in the ovaries of 14-day-old mice

S8 Video: Imaging of primordial follicles in human ovarian tissue

